# Elucidation of Codon Usage Signatures across the Domains of Life

**DOI:** 10.1101/421487

**Authors:** Eva Maria Novoa, Olivier Jaillon, Irwin Jungreis, Manolis Kellis

**Affiliations:** Computer Science and Artificial Intelligence Lab, MIT, Cambridge 02139, MA, USA; Broad Institute of MIT and Harvard, Cambridge 02139, MA, USA; Garvan Institute of Medical Research, Darlinghurst 2010 NSW, Australia; University of New South Wales Sydney, Sydney NSW Australia; Génomique Métabolique, Genoscope, Institut François Jacob, CEA, CNRS, Univ Evry, Université Paris-Saclay, 91057 Evry, France

**Author notes:** Corresponding authors: Eva Maria Novoa and Manolis Kellis.

## Abstract

Due to the degeneracy of the genetic code, multiple codons are translated into the same amino acid. Despite being ‘synonymous’, these codons are not equally used. Selective pressures are thought to drive the choice among synonymous codons within a genome, while GC content, which is generally attributed to mutational drift, is the major determinant of interspecies codon usage bias. Here we find that in addition to the bias caused by GC content, inter-species codon usage signatures can also be detected. More specifically, we show that a single amino acid, arginine, is the major contributor to codon usage bias differences across domains of life. We then exploit this finding, and show that the identified domain-specific codon bias signatures can be used to classify a given sequence into its corresponding domain with high accuracy. Considering that species belonging to the same domain share similar tRNA decoding strategies, we then wondered whether the inclusion of codon autocorrelation patterns might improve the classification performance of our algorithm. However, we find that autocorrelation patterns are not domain-specific, and surprisingly, are unrelated to tRNA reusage, in contrast to the common belief. Instead, our results reveal that codon autocorrelation patterns are a consequence of codon optimality throughout a sequence, where highly expressed genes display autocorrelated ‘optimal’ codons, whereas lowly expressed genes display autocorrelated ‘non-optimal’ codons.

## Introduction

Despite the relative universality of the genetic code and the conservation of the translation machinery across species, synonymous codons are not equally used, and codon biases vary dramatically between organisms and across genes within the same genome (Hershberg and Petrov 2008; Plotkin and Kudla 2011; Novoa and Ribas de Pouplana 2012; Shabalina, et al. 2013). Various factors can influence codon usage bias within and across genomes, including protein expression level (Gouy and Gautier 1982; Ikemura 1985), GC content (Hershberg and Petrov 2009; Palidwor, et al. 2010), recombination rates (Marais, et al. 2001), translation efficiency (Sorensen, et al. 1989; Tuller, Waldman, et al. 2010; Qian, et al. 2012), mRNA structure (Kudla, et al. 2009), codon position (Tuller, Carmi, et al. 2010), mRNA stability (Presnyak, et al. 2015) and gene length (Eyre-Walker 1996; Duret and Mouchiroud 1999), amongst others.

Although each species has a preference towards a specific subset of codons (Hershberg and Petrov 2009; Plotkin and Kudla 2011), the origin and evolutionary pressures driving these preferences remains largely unknown. Codon usage variation within genomes (intra-species codon usage) is often attributed to selection, due to the significant positive correlation between protein expression levels and the presence of “preferred” or “optimal” codons (Sharp, et al. 1986; Duret and Mouchiroud 1999). Indeed, codon bias is more extreme in highly expressed genes to match the skew of tRNA gene pools, providing a fitness advantage due to increased efficiency and/or accuracy in protein synthesis (Bulmer 1991; Akashi 1994; Dong, et al. 1996; Duret 2000; Gingold and Pilpel 2011). In contrast, the processes that drive codon usage variation across genomes (inter-species codon usage) are generally thought to be mutational (Chen, et al. 2004; Hershberg and Petrov 2008; Sharp, et al. 2010), although the extent to which these processes are driven by mutation, selection, or biased gene conversion remains controversial (Lassalle, et al. 2015; Long, et al. 2018). Genomic GC content has been identified as the strongest determinant of codon usage variation across species (Knight, et al. 2001; Palidwor, et al. 2010). Consequently, GC-rich organisms tend to favor GC-rich codons while AT-rich organisms are enriched in AT-rich codons (Hershberg and Petrov 2009).

tRNA gene content and codon usage bias are thought to coevolve (Dong, et al. 1996; Yona, et al. 2013) as a means to modulate translation speed for accurate co-translational protein folding (Komar 2009; Yu, et al. 2015). Indeed, tRNA deletions in *S. cerevisiae* are recurrently corrected, with the anticodon of a second tRNA mutated to match that of the deleted tRNA (Yona, et al. 2013). Supporting this, species belonging to the same domain of life (Archaea, Bacteria, Eukarya) evolved similarly in terms of their tRNA gene contents and decoding strategies (Novoa, et al. 2012), despite large differences in GC content. In the light of these observations, here we hypothesize and test whether species from the same domain of life may display similar codon usage biases despite their differences in GC content.

The elucidation of which evolutionary pressures have shaped extant genomes is crucial to comprehending why and how genomes evolve, but also can be exploited to build algorithms that can taxonomically annotate any given genomic sequence based on its properties. In this regard, next-generation sequencing has provided a great opportunity to explore complex ecological systems, such as microbiomes from the human gut or environmental samples. However, these samples often include a significant portion of uncharacterized species, and consequently, assigning sequence scaffolds to individual species, or even to higher-level taxa, remains challenging. Metagenomic annotation solutions, also known as ‘binning’, often rely on similarity-based searches (Brady and Salzberg 2009; Gerlach and Stoye 2011; Huson, et al. 2016). Unfortunately, such homology-based methods are unable to correctly annotate a significant portion of sequences (Prakash and Taylor 2012), such as those that are taxonomically-restricted, or do not have detectable homologues in other lineages. *De novo* taxonomical predictions, independent of homology, can be extremely useful for these situations.

Here we show that after removing variation associated with GC content, species from the same domain share similar codon bias signature, and identify that the codon usage bias of a single amino acid, arginine, is largely responsible for the separation of the species into their corresponding domains. We then show that coding sequences can be correctly classified into their corresponding domain of life, with an accuracy of 85%, using exclusively their codon usage biases. We speculate that domain-specific preferences for arginine codons are related to translation speed, which would support the view that codon usage variation across genomes is shaped not only by mutational biases, but also by selective forces.

## Results

### Beyond GC content, codon usage bias shows domain-specific patterns

The non-uniform usage of synonymous codons in a given sequence or genome can be measured as relative synonymous codon usage (RSCU), which is defined as the ratio of the observed frequency of codons to the expected frequency. In other words, the RSCU represents the deviation of the observed codon usage from a uniform distribution in which all codons encoding for the same amino acid have the same probability (Sharp, et al. 1986). Therefore, the codon usage bias of each species can be represented by a 59-dimensional RSCU vector (one per codon, removing Trp, Met, and stop codons). Upon hierarchical clustering of species based on their average RSCU, we find that species do not cluster following the tree of life, but rather, based on GC content, suggesting GC content is the major determinant of codon usage bias across species (**Figure 1A**), in agreement with previous works (Hershberg and Petrov 2009).

**Figure 1.**
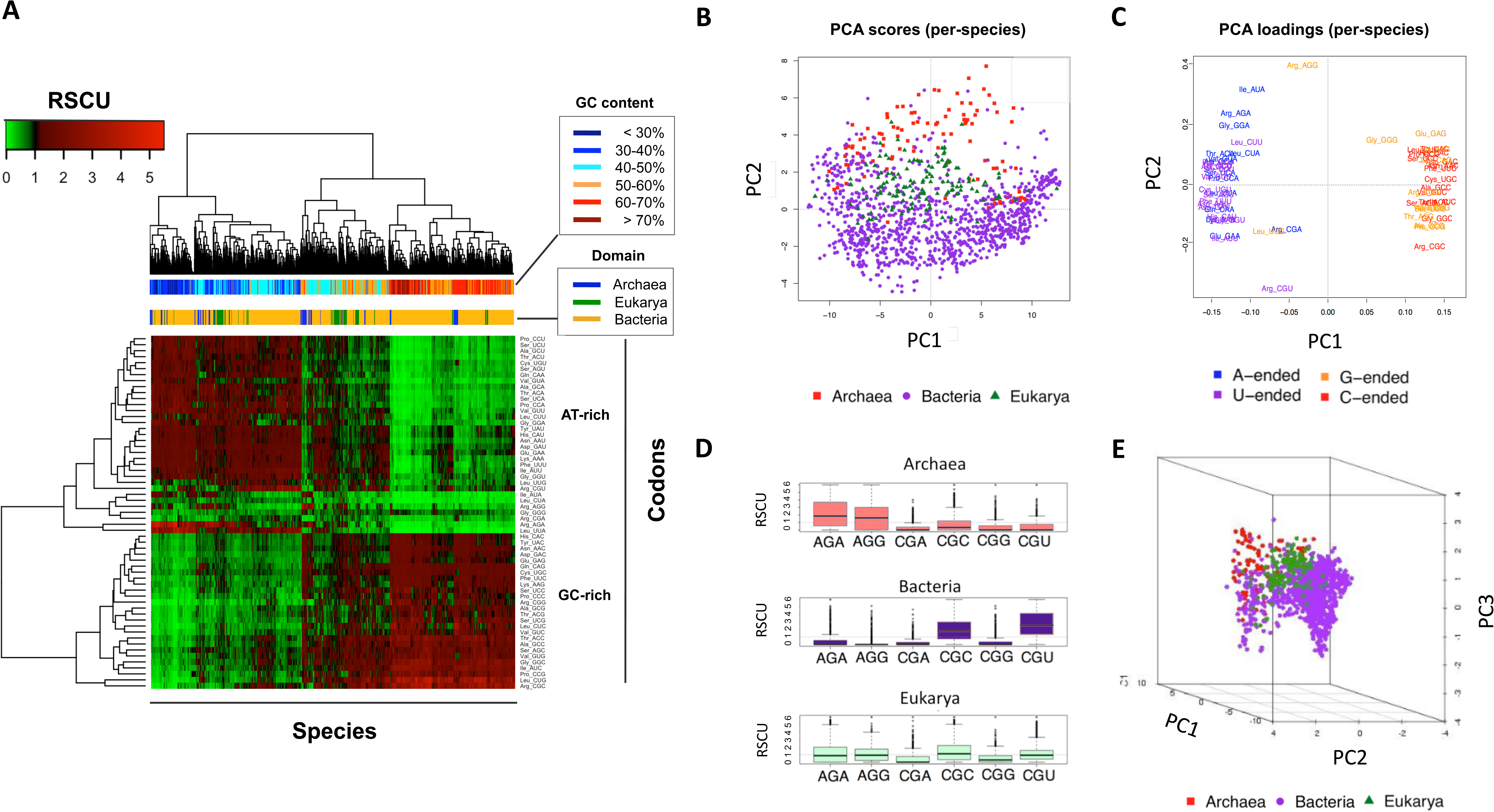
Analysis of codon usage bias across the three domains of life. (**A**) Hierarchical clustering of the average relative synonymous codon usage (RSCU) for each species (n=1765). Horizontal bars indicate GC content and domain of life for each species, and show that RSCU clusters species primarily by their GC content rather than by domain (**B**) Scatter plot of the first two principal components of the of the matrix of RSCU values in panel A. Each dot represents a species, and has been colored according to its corresponding domain. (**C**) PCA loadings plot for the first two principal components, where each codon has been coloured according to its ending nucleotide: G (orange), C (red), A (blue) or T (purple), showing that the PC1 score of a species is primarily determined by differences in frequencies of codons ending in GC or AT, whereas PC2 is mainly driven by differences in frequencies of arginine codons more than those of any other amino acid (**D**) Boxplot representation of arginine codon usage for each domain of life, showing that Archaea favor AGA and AGG codons, Bacteria favor CGC and CGU codons, and Eukarya show intermediate preferences. (**E**) 3D scatter plot representing each species by its first three principal component scores, using as input only the RSCU values of arginine codons, showing that arginine codon usage alone allows for discrimination of domains. See also Figure S1.

To deconvolute the bias related to GC content from that caused by codon usage, we applied principal component analysis (PCA) to the average RSCU of each analyzed species (see Methods), with the expectation that the first principal component (PC1) would capture the variance due to GC content. Indeed, we find that PC1 does not separate the species into domains (**Figure 1B**, see also **Figure S1**), but clusters them according to GC content, as reflected by the contributions of the individual codons to PC1 (i.e. GC-ended codons have negative PC1 scores, whereas AT-ended codons have positive PC1 scores) (**Figure 1C**). In contrast, the second principal component (PC2) is capable of separating species into their corresponding domains of life (**Figure 1B**), confirming our hypothesis.

### Arginine codons are the major drivers of inter-species codon usage bias

Two codons, CGU and AGG, both coding for arginine, are the largest contributors to the separation of species in PC2 (**Figure 1C**). Upon closer examination of arginine codon usage biases in the three domains, we observe that their relative usage of arginine codons is distinct (**Figure 1D**). More specifically, Archaea preferentially use AGG and AGA, whereas Bacteria preferentially use CGC and CGU, and Eukarya show intermediate preferences between Archaea and Bacteria.

We then wondered whether the arginine codon bias was sufficient to cluster species into their corresponding domains. To test this, we repeated the PCA analysis as described above, but this time using exclusively arginine codon biases, finding that the differences in arginine codon usage across species were sufficient to recapitulate the clustering of species into domains (**Figure 1E**). Therefore, we conclude that arginine codon usage bias is a major contributor to inter-species codon usage bias across domains.

### Arginine codon usage bias is relatively stable within species

Codon usage bias varies within genomes, with highly expressed proteins more enriched in ‘optimal’ of ‘preferred’ codons, compared to proteins with lower expression (Bulmer 1987; Duret 2000; Higgs and Ran 2008; McDonald, et al. 2015). Therefore, we wondered whether our findings would be applicable at the level of individual sequences. For this aim, we individually analyzed all sequences from the 1765 EMBLCDS species from all three domains (see Methods). We computed RSCU values for each sequence, performed PCA dimensionality reduction, and retained the first 3 principal components for further analysis, based on Scree’s test (Cattell 1966). We found that individual sequences also clustered by domain, suggesting that intra-species codon usage variation is not larger than inter-species codon usage variation (**Figure 2**).

**Figure 2.**
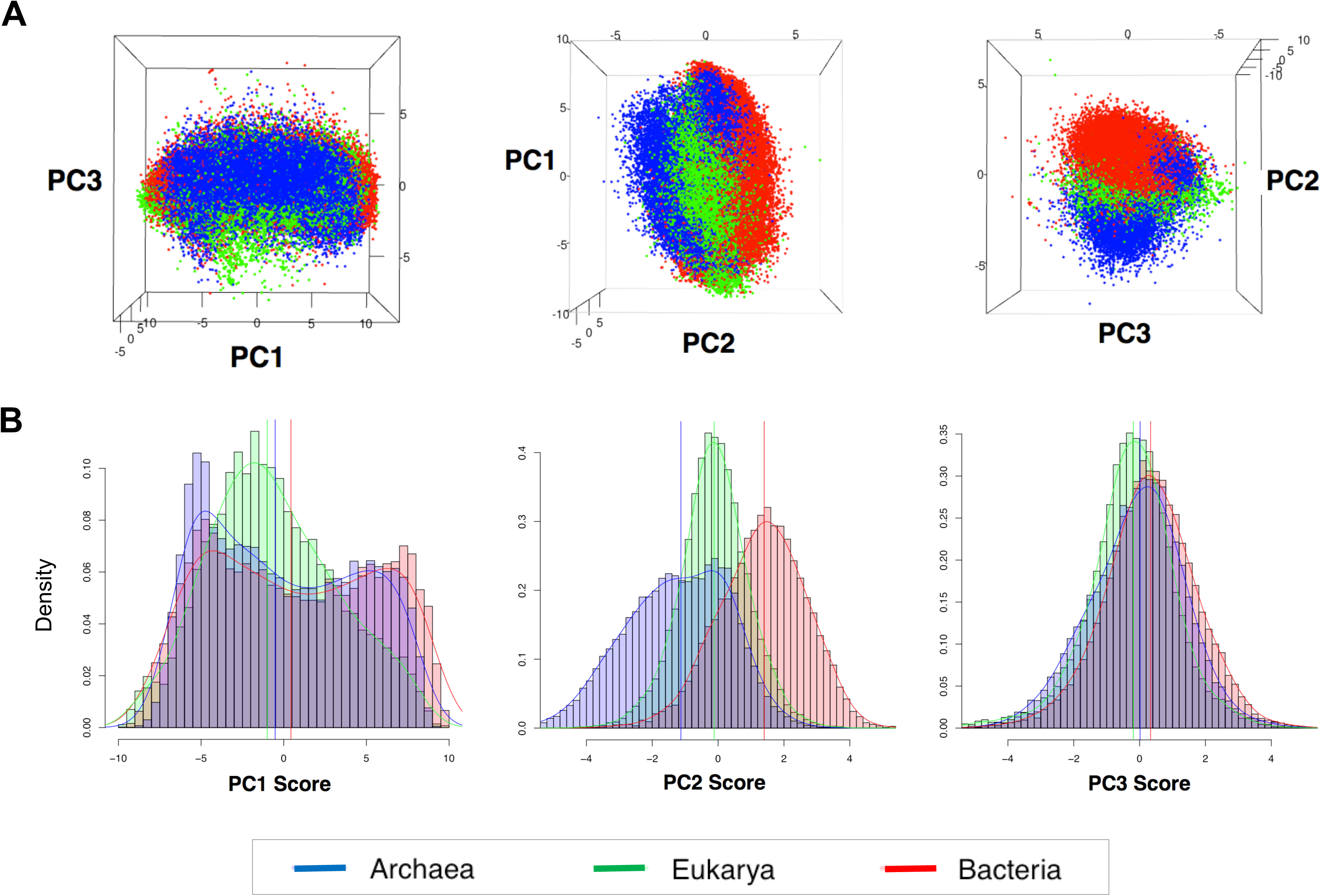
Codon usage bias clusters sequences into their corresponding domains. (**A**) 3D scatter plot of the first three principal component scores for all EMBLCDS sequences included in the analysis. Each dot represents a sequence, and has been coloured by its corresponding domain of life: Archaea (blue), Bacteria (red), Eukarya (green). (**B**) Histograms of the densities of the PC scores for each domain: PC1 scores (left), PC2 scores (middle), PC3 scores (right).

Surprised by these results, we hypothesized that although global codon usage may strongly vary between highly and lowly expressed genes (Bulmer 1987; Duret 2000; Higgs and Ran 2008; McDonald, et al. 2015), this might not be the case for all amino acids, such as arginine. To test this, we examined the codon usage of all coding sequences (CDS) of *S. cerevisiae*, and determined how codon usage varied with protein expression –using previously published proteomics datasets (Newman, et al. 2006)– for each individual amino acid and codon subtype (**Figure 3**). As expected, we observed that codon usage drastically varied depending on protein abundance. For the majority of amino acids, codon preferences completely switch from ‘non-optimal’ to ‘optimal’ depending on expression level (Lys, Asn, His, Phe, Asp, Tyr, Ala, Thr, Val, Ser, Ile). However, in other cases, codon preferences are maintained –although become more extreme– when comparing lowly and highly expressed proteins (Gln, Glu, Cys, Gly, Arg). The relative consistency of arginine codon usage across a genome explains why our analyses can be applied not only at an average per-species level, but also at the level of individual sequences.

**Figure 3.**
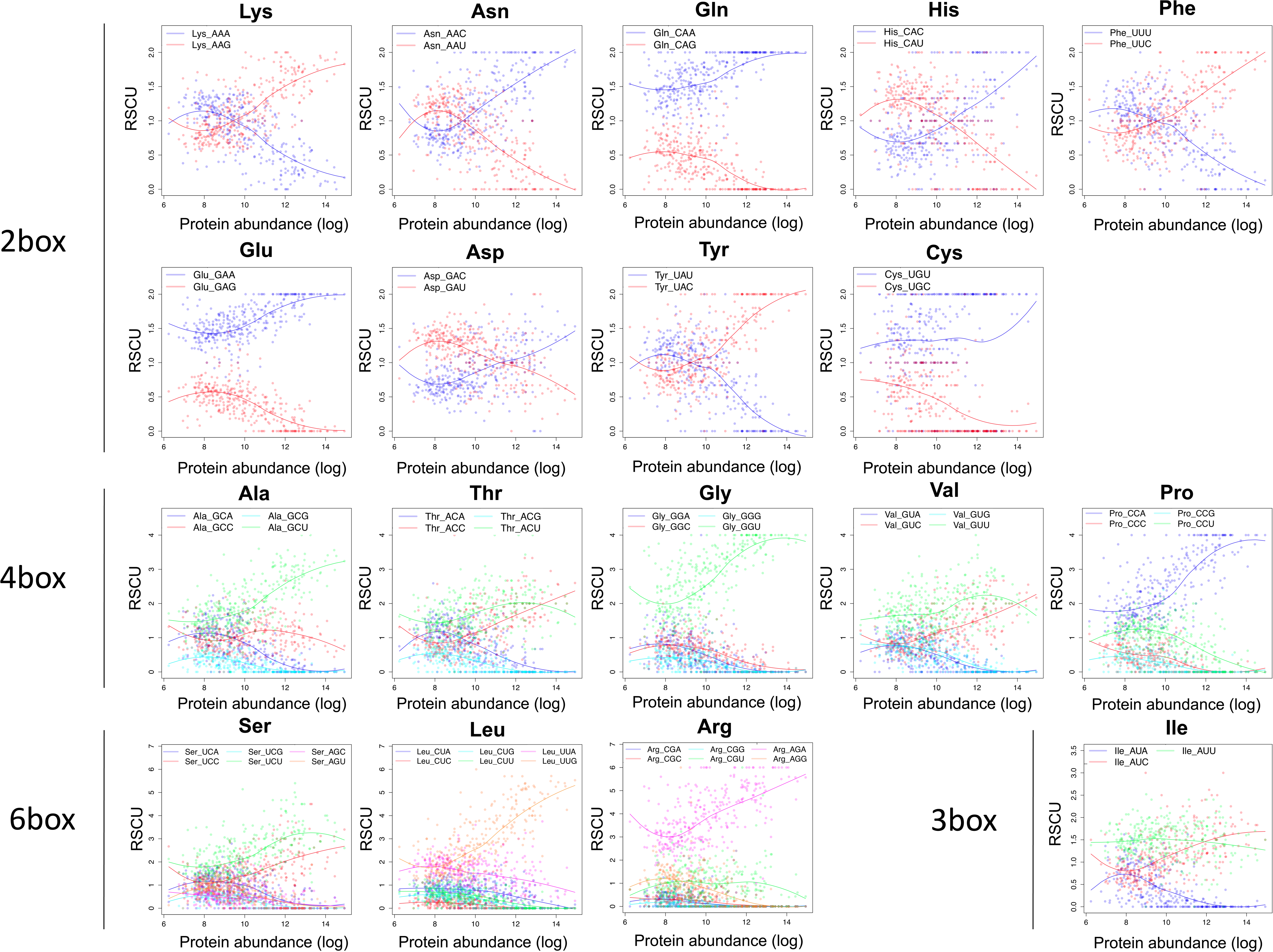
Codon preferences in *S. cerevisiae* as a function of expression levels. Codon preferences, represented by relative synonymous codon usage (RSCU), are reversed between highly and lowly expressed genes for some amino acids but not for others. Although codon usage varies within a genome, intra-genome differences are small enough that individual sequences still cluster by domain, as seen in Figure 2.

### Codon usage signatures can be used to taxonomically annotate orphan sequences from metagenomic analyses

We then wondered whether simple patterns of codon usage bias would be sufficient to classify a species into its corresponding domain of life. To test this, we used the previously built 59-dimensional RSCU vectors for each EMBLCDS sequence, subdivided the data into training and testing sets, and built a Support Vector Machine (SVM) with the training set data (**Figure 4A**). We find that codon usage bias alone predicts the correct domain with AUC values ranging from 0.78-0.84 (**Figure 4B**). The accuracy of prediction is dependent on the sequence length as could be expected (**Figure S2**), however, predictions were found to be better than random even when analysing the shortest set of CDS sequences (100-200 nt).

**Figure 4.**
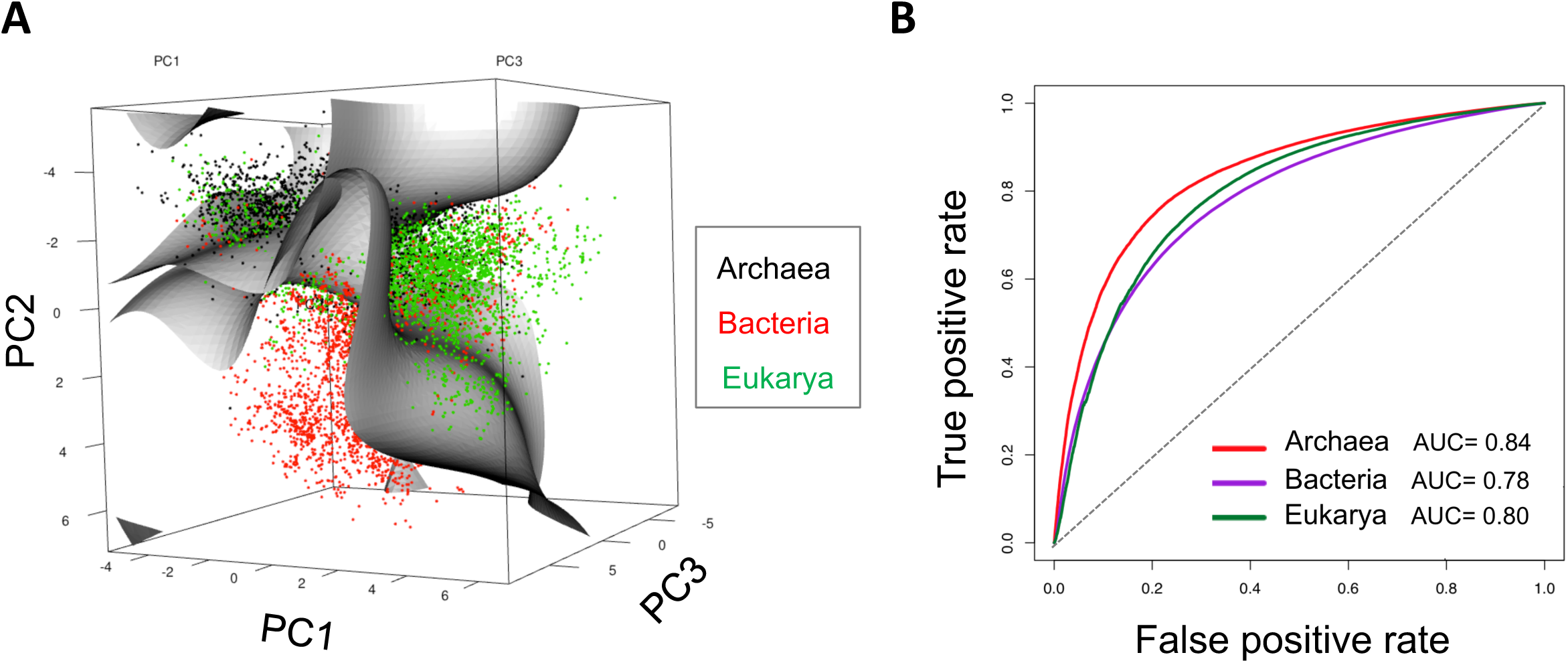
Taxonomical classification of sequences using codon usage bias. (**A**) 3D plot representation of the first three principal component scores. Support Vector Machine hyperplanes computed using the training set are also shown. Each dot represents a sequence, and has been coloured according to its corresponding domain of life. (**B**) ROC curves of the SVM class probabilities, computed separately for each domain. See also Figure S2.

### Codon autocorrelation does not reflect tRNA reuse

Previous studies in yeast have shown that once a particular codon has been used, subsequent occurrences of the same amino acid in the same transcript are not random (Cannarozzi, et al. 2010), a phenomenon termed as ‘codon autocorrelation’ or ‘codon covariation’. Mechanistically, it was argued that tRNA recycling was the driving force causing the observed biased distribution of synonymous codons along a sequence, i.e., codons that would reuse the same tRNA would be favored as a means to increase the speed of translation (Cannarozzi, et al. 2010). A subsequent study re-examined this question, and compared the autocorrelation between codons encoding the same amino acids to those encoding different ones (Hussmann and Press 2014). Intriguingly, this second study found that covariation between codons encoding different amino acids was as strong as covariation between codons encoding the same amino acid, concluding that there was insufficient evidence to claim that tRNA recycling is the force driving codon autocorrelation. Despite the uncertain cause of codon covariation, both studies show that the probability of observing a specific codon is dependent on previous codon occurrences, at least in the case of *S. cerevisiae.*

Considering that species from the same domain of life share common codon usage signatures (**Figure 1B-E** and **Figure 2**), we wondered whether codon covariation would also follow a similar behavior. For this aim, we calculated codon covariation as described in Cannarozzi et al. (Cannarozzi, et al. 2010) (see Methods), finding that codon covariation within same amino acids in *S. cerevisiae* partly supports a tRNA recycling model (**Figure 5A**). For example, in the case of alanine codons, GCA and GCG show covariation, and are both decoded by tRNA^Ala^_UGC_. Similarly, GCC and GCT are decoded by tRNA^Ala^_AGC_ and also show covariation, in agreement with a tRNA recycling model. In contrast, the covariation observed in other species was not supported by a tRNA recycling model (**Figure 5B-D**). Taking as an example the same amino acid, alanine, covariation in human sequences was detected between GCA and GCT codons, which are decoded by two different tRNAs, tRNA^Ala^_UGC_ and tRNA^Ala^_AGC_, as well as between GCG and GCC codons, despite being decoded by two different tRNAs, tRNA^Ala^_CGC_ and tRNA^Ala^_AGC_ (**Figure 5B**). Similarly, in *E. coli*, the two alanine codons that show covariation are GCA and GCT, despite being decoded by two different tRNAs, tRNA^Ala^_UGC_ and tRNA^Ala^_GGC_ (**Figure 5C**). Overall, our results suggest that codon covariation is unrelated to tRNA reusage, in agreement with the second study described above (Hussmann and Press 2014).

**Figure 5.**
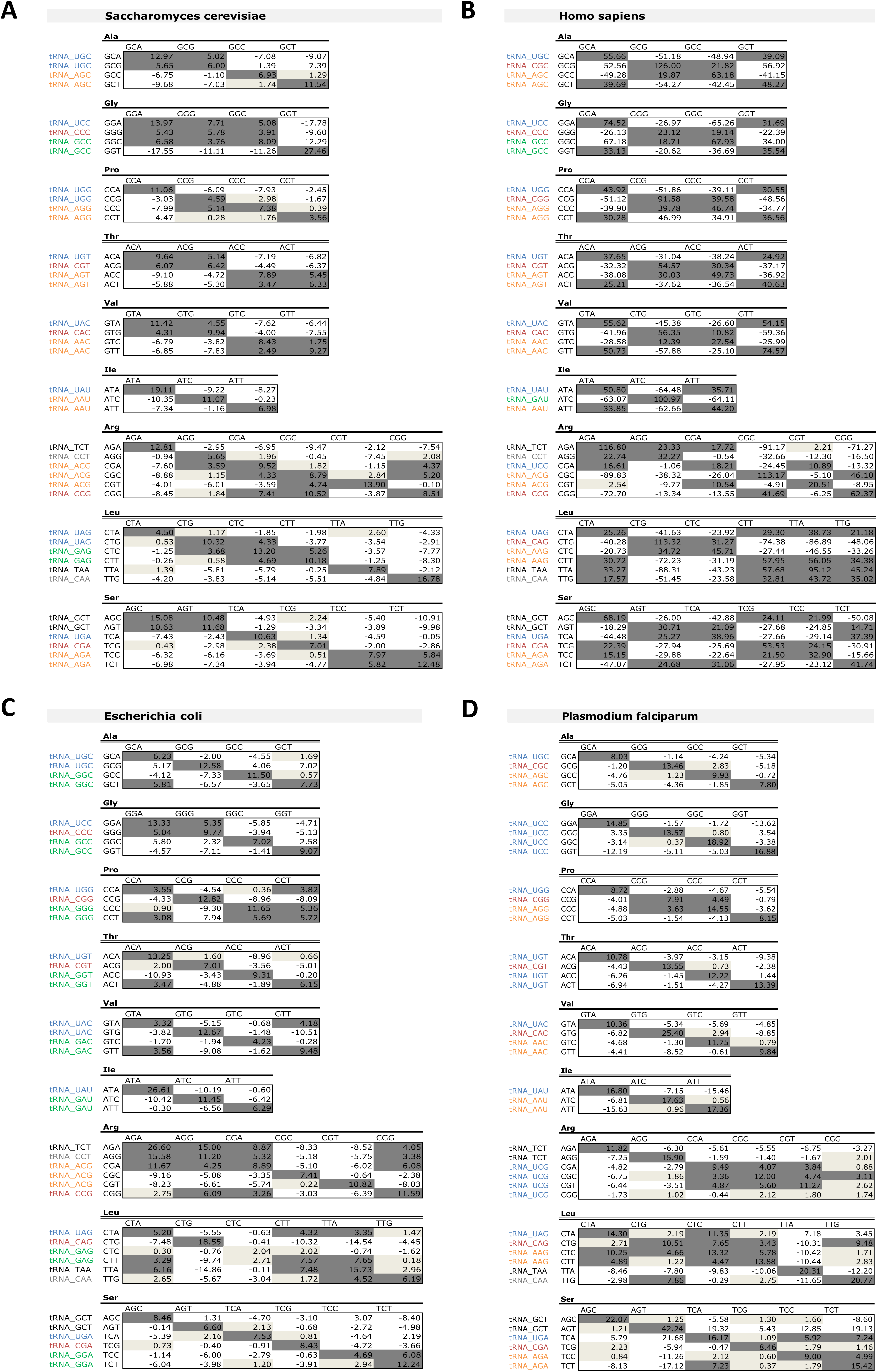
Analysis of codon covariation across species does not support a universal tRNA recycling model. Codon covariation measured over all pairs comprised of one codon and the subsequent one encoding for the same amino acid, shown for *S cerevisiae* (**A**), *H. sapiens* (**B**), *E. coli* (**C**) and *P. falciparum* (**D**). Values correspond to standard deviations from expected. Each codon has been labelled with its corresponding decoding tRNA, following parsimony-extended wobble rules when no Watson-Crick matching tRNA isoacceptor is available (as per gtRNAdb (Chan and Lowe 2016)). Pairs have been shaded according to the number of standard deviations from expected: dark grey (>+3SD; strongly favoured codon pair), light grey (0-3SD; slightly favoured codon pair), white (<= 0 SD; non-favoured codon pair). In yeast, most codon pairs using the same tRNA are overrepresented, supporting a tRNA recycling model to explain the overrepresentation, but that is not true in other species.

### Codon autocorrelation reflects global sequence codon optimality

We then wondered whether codon covariations may be in fact a simple consequence of codon optimality throughout a sequence, i.e. whether ‘optimal’ codons, which are abundant in highly expressed proteins, would appear as autocorrelated, and ‘non-optimal’ codons, which are abundant in lowly expressed proteins, would also appear as autocorrelated. To test this, we compared the observed codon covariations in *S. cerevisiae* and *E. coli* with the set of ‘optimal’ and ‘non-optimal’ codons, defined as those that were highly or lowly abundant in highly expressed proteins, respectively (**Figure 6**). We find that after binning codons into ‘optimal’ and ‘non-optimal’, codon covariations were present within ‘optimal’ or within ‘nonoptimal’ codons, but not across them. More specifically, 97% (31/32) of the autocorrelated codon pairs (SD ≥3) in *S. cerevisiae* (**Figure 6A**) and 86% (25/29) of the autocorrelated pairs (SD ≥3) in *E. coli* (**Figure 6B**) could be explained by codon optimality. It is important to note that the remaining autocorrelated codon pairs in *E. coli* (4/29) and in S. cerevisiae (1/32) actually correspond to codons which we labeled as ‘intermediate optimal’ (yellow boxes), for which we considered that we could not clearly assign the category of ‘optimal’ or ‘nonoptimal’, and thus were not counted as positive results. Overall, our results suggest that, at least in the case of *S. cerevisiae* and *E. coli*, codon autocorrelation may be a consequence of similar choice of ‘optimal’ or ‘non-optimal’ codons throughout a sequence.

**Figure 6.**
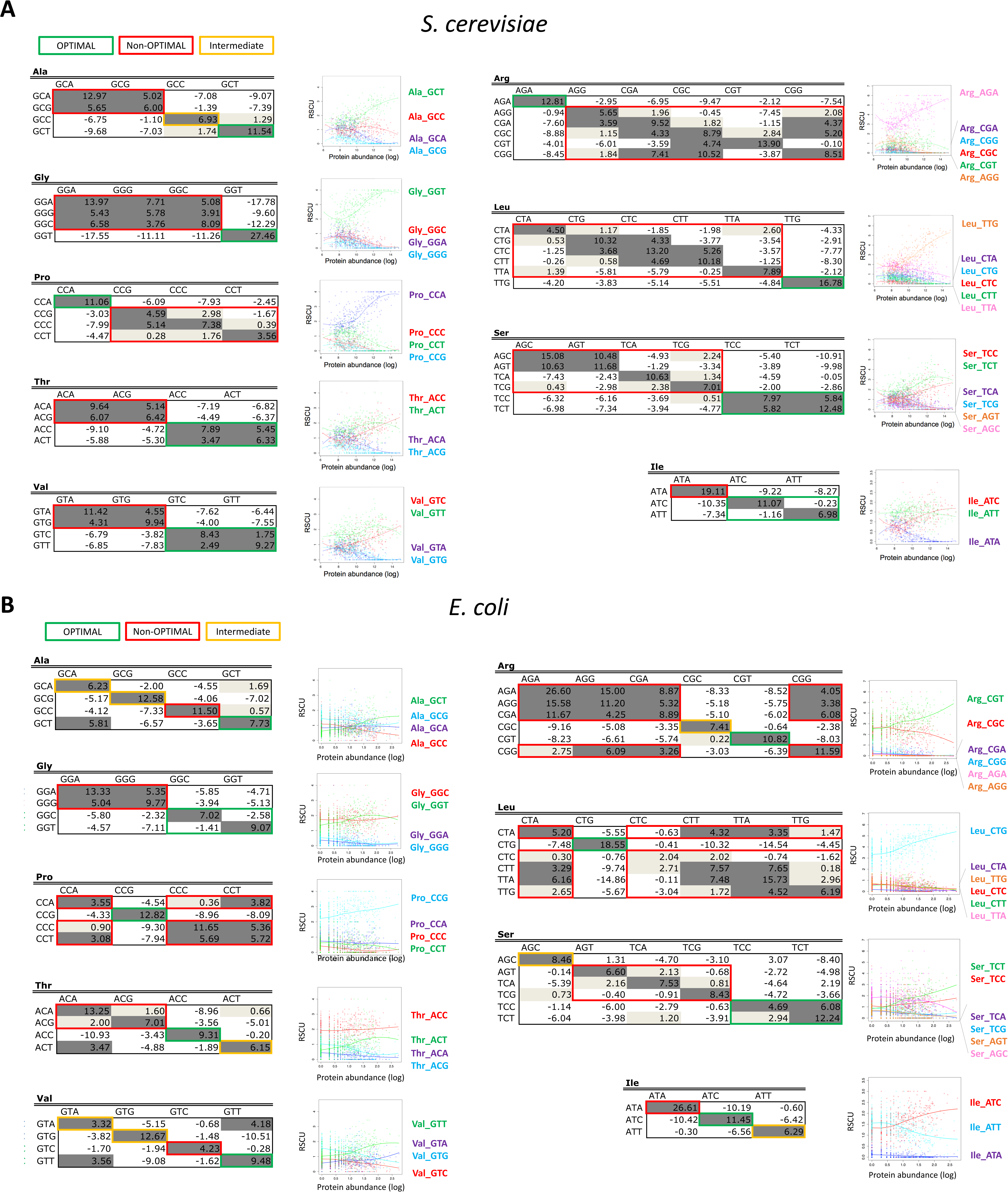
Codon covariation is likely a consequence of the co-occurrence of ‘optimal’ and ‘non-optimal’ codons, in highly and lowly expressed proteins, respectively. Codon covariation for *S. cerevisiae* (**A**) and *E. coli* (**B**) as depicted as in Figure 5, highlighting those pairs that are formed by two optimal codons (dark green), two non-optimal codons (red) and codons with intermediate optimality (yellow). Optimal and non-optimal codons have been defined as those that are highly abundant and lowly abundant in highly expressed proteins, and their relative abundance is shown for each individual amino acid and codon.

### Codon autocorrelation is not domain-specific

Regardless of the evolutionary forces driving codon covariation, our analysis demonstrates that codon covariation exists in all species analyzed (**Figure 5**). Therefore, we extended this analysis to hundreds of species across the 3 domains of life (see Methods), and examined whether taxonomically-related species displayed similar codon covariation patterns. We found that the number of standard deviations from the expected codon pair usage (Cannarozzi, et al. 2010) was not a useful metric to compare species, due to dependence of this metric on genome size (**Figure 5**). Therefore, we defined a new metric, independent of genome size, which we termed Relative Synonymous Codon Pair Usage (RSCPU) (see Methods). This metric represents the ratio of the observed usage of a given codon pair to the expected pair usage, which is defined as the product of the observed usage of the two individual codons in the genome. It is important to note that RSCPU values are normalized by the individual usage of the two codons in the genome, and thus are independent of GC content.

Our results show that taxonomically-related species display similar codon variation, however, multiple clusters appear within each domain of life, suggesting that codon covariation is not domain-specific (**Figure 7**). Nevertheless, we do observe that some species belonging to the same domain cluster together, suggesting that codon covariation is not completely independent of their taxonomy. We then performed PCA analysis on the RSCPU values to test whether additional principal components might better separate the species into their corresponding domains; however, we find that species belonging to different domains largely overlap (**Figure S3**). Overall, our results suggest that codon covariation patterns are not domain-specific, but they do show certain degree of clustering which is dependent on their taxonomy. Therefore, we suggest that codon covariation signatures may be used as additional features to improve the performance of current binning algorithms, but cannot be used alone to classify species into their corresponding domains of life.

**Figure 7.**
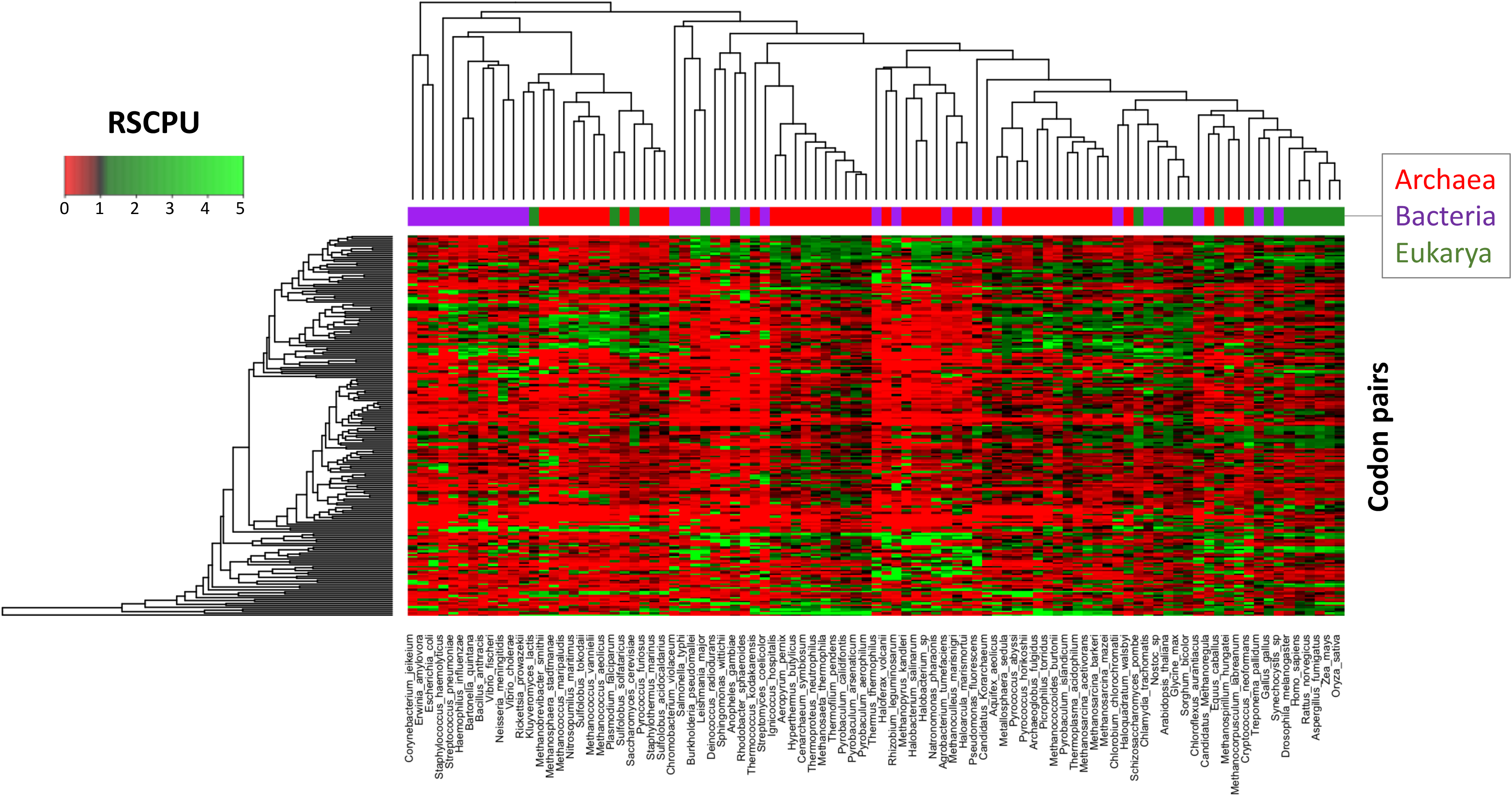
Hierarchical clustering of codon covariation patterns across species spanning the three domains of life. Each codon pair has been coloured according to its RSCPU value. The upper bar over the heatmap represents the corresponding domain of each species. See also Figure S3.

## Discussion

It is well established that the identity of favored codons varies among organisms (Chen, et al. 2004; Hershberg and Petrov 2009; Sharp, et al. 2010). However, the rules governing the identities of favored codons across organisms still remain obscure. High GC content organisms tend to have GC-rich favored codons, while low GC content organisms favor AT-rich codons, suggesting evolutionary pressures act in the same direction as the nucleotide substitution biases that determine overall nucleotide content of genomes (Hershberg and Petrov 2009). Based on these observations, previous works suggested that inter-species codon bias is driven mostly by genome-wide mutational biases. Here we suggest that, in addition to mutational forces, inter-species codon bias might also be shaped by selective forces.

Several studies have reported codon translation rates for a wide variety of species and conditions (Li, et al. 2012; Gardin, et al. 2014; Lareau, et al. 2014; Bazzini, et al. 2016). In a scenario of high abundance of nutrients, arginine codons have been recurrently identified as the slowest translated codons, both in Bacteria (Bonekamp and Jensen 1988; Chevance, et al. 2014) and Eukarya (Charneski and Hurst 2013; Gardin, et al. 2014; Requiao, et al. 2016). However, within a species, not all arginine codons display slow translation rates. For example, in a bacterial system based on *Salmonella enterica*, CGC codons are rapidly translated, whereas AGG codons, which also encode for arginine, are slowly translated (Chevance, et al. 2014). In the eukaryote *S. cerevisiae*, codons that are rapidly and slowly translated differ from those identified in bacteria. More specifically, CGC is slowly translated in *S. cerevisiae*, whilst AGA and CGU are rapidly translated (Gardin, et al. 2014), matching the codon preferences of *S. cerevisiae* (**Figure 3**). Moreover, in the case of Bacteria, the variance in translation speed between codons that encode for arginine is larger than for any other amino acid, and it is also high in the case of Eukarya (**Figure S4A**). In the light of these observations, we suggest that selective pressures towards maintaining specific arginine codon usage biases, compared to other amino acids, might be responsible for the existence of domain-specific arginine codon usage biases.

A remaining question, however, is why such extreme variance in translation speed of arginine codons exists. It has been suggested that the rare usage and slow translation speed of AGG codons in Bacteria is a consequence of their similarity to Shine-Dalgarno sequences (Li, et al. 2012; Chevance, et al. 2014) (**Figure S4B**). In agreement with this hypothesis, Shine-Dalgarno-like sequences are typically depleted in bacterial genomes (Li, et al. 2012). However, Shine-Dalgarno sequences are also employed by archaeal genomes to help recruit the ribosome to initiate protein synthesis, and in this domain, AGG codons –together with AGA codons– are in fact the most frequently used codons. Therefore, it is unlikely that the similarity to Shine-Dalgarno sequences alone is responsible for the depletion of AGG codons in Bacteria. An alternate –but not mutually exclusive– explanation for this phenomenon might reside in the differences between arginine tRNA decoding strategies used in Archaea and Bacteria (Novoa, et al. 2012). More specifically, in Bacteria, the appearance of tRNA adenosine deaminases (*tadA*), which allows for efficient translation of CGC and CGU codons, might explain why these two codons are more frequently used and rapidly translated in Bacteria (**Figure S4C**). Moreover, *tadA* does not exist in Archaea, and thus it may explain why archaeal genomes do not preferentially use CGN codons. Future work will be needed to decipher which are the forces causing distinct arginine codon usage bias across different domains of life.

Regardless of the evolutionary forces driving domain-specific codon preferences, here we find that codon usage bias of individual sequences can be used to taxonomically annotated at the domain level (**Figure 4**), largely due to differences in arginine codon usage (**Figure 1**). To improve the performance of the algorithm, we wondered whether we could also include codon covariation, which is a well-documented phenomenon in yeast (Cannarozzi, et al. 2010; Hussmann and Press 2014). We find complex covariations of codon pairs are present in all analyzed species (**Figure 5**), which do not seem to be related to tRNA recycling, but rather, to similar codon optimality throughout a sequence (**Figure 6**). Unfortunately, codon covariation patterns were not shared by the species within each domain of life (**Figure 7** and **S3**).

Composition-based binning methods have been previously applied to taxonomically annotate metagenomic sequences (McHardy, et al. 2007; Brady and Salzberg 2009; Diaz, et al. 2009; Rosen, et al. 2011; Alneberg, et al. 2014; Lin and Liao 2016; Lu, et al. 2017). These methods exploit the uniqueness of base composition – from single to oligonucleotide levels – found across the genomes of different taxonomic entities, and have been implemented in tools such as PhyloPhythia (McHardy, et al. 2007), TETRA (Teeling, et al. 2004) and TACOA (Diaz, et al. 2009), amongst many others. However, the underlying evolutionary principles governing the observed k-mer variation across species remains largely unknown. Consequently, it may be difficult to improve such algorithms without a working hypothesis of which additional variables might affect performance. Here we find that taxonomically-related species display common covariation patterns, although these patterns are not domain-specific. In the light of these findings, we propose that in addition to considering features such as overall k-mer counts (Teeling, et al. 2004; Lu, et al. 2017), codon covariation may be used to further improve the performance of current composition-based metagenomics binning algorithms.

## Materials and Methods

### Gene sequences

The full set of EMBL CDS sequences was downloaded in July 2015 from ftp://ftp.ebi.ac.uk/. To avoid over-represented subsets of sequences, we selected those sequences that belonged to genomes with more than 1000 genes in the dataset, and one strain per species. The final set of EMBL CDS sequences analyzed consisted in 1765 genomes, which included over 11 million sequences.

### Training and test set preparation

The filtered set of EMBL CDS sequences was divided into training (10%) and test set (90%). Each training set sequence was individually analyzed in terms of codon usage bias, and converted into a 59-element vector (one for each codon, excluding Met, Trp, and stop codons) of relative synonymous codon usage values (RSCU). RSCU was computed as defined in Sharp et al. (Sharp, et al. 1986). Principal component analysis (PCA) was applied on the training set matrix of RSCU values to reduce the dimensionality of the data. Scree’s test (Cattell 1966) was used to determine the number of significant principal components to be retained. PCA scores of the selected subset of principal components were used as new vectors to define each sequence. Each sequence was assigned to a domain (Eukarya, Bacteria, Archaea) based on its taxonomical annotation in NCBI taxonomy.

### Machine learning

PCA scores of the EMBL CDS training set with its corresponding domain annotations were used as input to train a Support Vector Machine (SVM) using the e1071 library from R, using a C-classification method. Parameters were optimized using the tune.svm function. The final SVM model was validated using 5-fold cross-validation. EMBL CDS test set sequences were converted into PCA scores by applying the same PCA loadings that were generated upon PC analysis on the training set, and its corresponding domains were predicted using the SVM model built on the training set. Our model correctly predicted the domain of the EMBL CDS test set sequences with an overall accuracy of 85%.

### Codon autocorrelation

Codon autocorrelation standard deviations were computed as described in Cannarozzi et al (Cannarozzi, et al. 2010). Briefly, for each sequence, the number of consecutive pairs of codons for a same amino acid were counted. The expected numbers of pairs were computed as the products of the frequencies of the individual codons. Codons were Z-transformed by subtracting the expected counts from the observed and divided by the standard deviations from the expected value. The results were expressed as standard deviations from the expected value.

### Relative synonymous codon pair usage

We define the relative synonymous codon pair usage (RSCPU) as the ratio of the observed frequency (f_obs_pair_) of a given codon pair to the expected frequency of the codon pair (f_exp_pair_) (Eq.1). The expected frequency of the codon pair is defined as the product of the individual codon frequencies (f_obs_codon1_ and f_obs_codon2_) observed in the genome (Eq. 2).

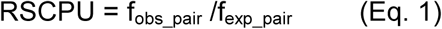

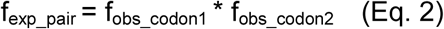

Therefore, if the observed frequency matches the expected frequency, the RSCPU will have a value of 1. If the RSCPU is higher than 1, the pair is seen more frequently than what would be expected considering the individual codon frequencies, whereas if it is lower than 1, the pair is observed less frequently than what would be expected.

### Protein abundances

Protein abundance values of *S. cerevisiae*, used to build Figure 3 and 6, were taken from the work of Newman et al. 2006 (Newman, et al. 2006). Protein abundance values of *E. coli*, used to build Figure 6, were taken from Lu et al. 2007 (Lu, et al. 2007).

## Acknowledgements

EMN was supported by a long-term postdoctoral fellowship from the Human Frontier Science Program (LT000307/2013-L), and is currently supported by a Discovery Early Career Researcher Award (DE170100506) from the Australian Research Council. IJ was supported by R01 HG004037 and GENCODE Wellcome Trust grant U41 HG007234 We thank all members of the Kellis lab for their valuable comments and suggestions.

## Supplementary Figure Legends

**Figure S1. (A)** Scatter plot of the first four principal component scores of the per-species average RSCU values. Each dot represents a species, and has been colored according to its corresponding domain of life. (**B**) Score density plots of the first four principal components, for each domain of life: Archaea (blue), Bacteria (red), Eukarya (green).

**Figure S2.** ROC curves of the SVM class probabilities, where CDS sequences have been binned based on their sequence length, showing that classification improves with sequence length, as expected. CDS sequences longer than 700 nucleotides were not included in the analysis.

**Figure S3**. Scatter plot of the first two principal components (left panel), and second and third components (right panel) of the scores of the matrix of RSCPU values of the 1765 EMBLCDS species used in this work show that species belonging to different domains largely overlap. Each dot represents a species, and has been colored according to its corresponding domain.

**Figure S4. (A)** Variance in translation speed between codons that encode for the same amino acid, using the data from Chevance et al. for a bacterial system (Chevance, et al. 2014) and the data from Gardin et al. for an eukaryotic system (Gardin, et al. 2014). Amino acids have been sorted according to their variance in Bacteria, and normalized to the amino acid with highest variance, for each domain of life separately. (**B**) Schematic representation of the pairing of the Shine-Dalgarno sequence upstream the start codon with the 16s rRNA, promoting ribosome recruitment. (**C**) Representation of the tRNA:codon pairings that occur for decoding arginine codons in Archaea and Bacteria. In the upper panels, the average relative synonymous codon usage (RSCU) values for each arginine codon are depicted. In the bottom panels, the average relative gene frequencies (RGF) for each tRNA arginine isodecoder are depicted. In Archaea, tRNA^Arg^(ACG) isoacceptors do not exist. In Bacteria, tRNA^Arg^(ACG) is converted to tRNA^Arg^(ICG) by tRNA adenosine deaminases (tadA), which can then decode CGC, CGU and CGA codons. It is worth noticing that inosine preferentially pairs with C and U (i.e. CGC and CGU codons in the case of arginine). The appearance of tadA enzymes in Bacteria might be responsible for the preferred usage of CGC and CGU arginine codons in Bacteria, whereas the lack of this enzyme in Archaea might explain why AGA and AGG codons are preferentially used in this domain.

